# Reproducible data management and analysis using R

**DOI:** 10.1101/644625

**Authors:** Bjørn Fjukstad, Nikita Shvetsov, Therese H. Nøst, Hege Bøvelstad, Till Halbach, Einar Holsbø, Knut Hansen, Eiliv Lund, Lars Ailo Bongo

## Abstract

**Background:** Standardizing and documenting computational analyses are necessary to ensure reproducible results. It is especially important for large and complex projects where data collection, analysis, and interpretation may span decades. Our objective is therefore to provide methods, tools, and best practice guidelines adapted for analyses in epidemiological studies that use -omics data.

**Results:** We describe an R-based implementation of data management and preprocessing. The method is well-integrated with the analysis tools typically used for statistical analysis of -omics data. We document all datasets thoroughly and use version control to track changes to both datasets and code over time. We provide a web application to perform the standardized preprocessing steps for gene expression datasets. We provide best practices for reporting data analysis results and sharing analyses.

**Conclusion:** We have used these tools to organize data storage and documentation, and to standardize the analysis of gene expression data, in the Norwegian Women and Cancer (NOWAC) system epidemiology study. We believe our approach and lessons learned are applicable to analyses in other large and complex epidemiology projects.

## Introduction

Reproducibility is necessary to advance science and to leverage scientific results [1]. This requires implementing best practices for data management and analysis. Such best practices are also necessary for large and complex projects where data collection, analysis, and interpretation may span decades, and is therefore done in several iterations, by different people. We have observed this need in systems epidemiology [2]. In addition, the need is recognized in the STROBE-ME [3] initiative to strengthen the reporting of observational studies in molecular epidemiology.

There are many approaches, systems, and tools for data storage and processing that solve many of the technical challenges of ensuring reproducible analyses [4]. To make it easy to find relevant data for re-analysis or re-interpretation, the data can be organized in file system structures, databases, or in other indexable storage systems. To keep track of different versions of files, we can can use a versioned file system, or version control systems such as *git*, that are widely adopted in software engineering. To document the tools, parameters, and reference databases used in an analysis we can use frameworks such as CWL [5], Galaxy [6], Snakemake [7], Spark [8], or an in-house solution such as our *walrus* system [9]. All of these frameworks provide an interface to set up an analysis pipeline, either as a text file or using a Graphical User Interface (GUI), and then execute it. To record provenance and keep track of the intermediate files, we can implement and run the analysis in for example Galaxy, Spark or Pachyderm [10]. However, there is a need to adapt these systems and tools to the needs of typical omics data analysis workflows.

In this paper we describe our lessons learned in 10 years of transcriptomics data analysis in the Norwegian Women and Cancer (NOWAC) study [11]. We use these to propose an approach to maintain, preprocess, and facilitate statistical analyses in complex systems’ epidemiology datasets. The approach ensures reproducibility, and we believe that it is well adapted for omics data analyses. It enables us to achieve reproducible research through the four steps described above. First, we use R since it has many up-to-date and actively maintained packages for analyzing, plotting, and interpreting data (for instance, Bioconductor [12] and the Comprehensive R Archive Network [13]). Second, we have developed an R package with code and the datasets from the NOWAC study. We document all datasets thoroughly and use version control to track both datasets and code over time. Third, we have developed an interactive web application, the *Pippeline*, to perform the standardized preprocessing steps for gene expression datasets. Fourth, we export the data as a git repository and RStudio project file to encourage reproducible analyses. Fifth, we have developed our own best practices to report results and share analyses through reproducible analysis reports.

The article is organized as follows. After the description of the datasets at hand and given context, we detail how omics data analysis was done previously, and what challenges this implied. We then discuss the requirements for our new approach and describe the solution in detail, together with an explanation of the corresponding methodology and best practices. We briefly discuss limitations of our work before the concluding.

## Data analysis lessons learned in the Norwegian Women and Cancer study

Our approach is based on the 10 years of transcriptomics data analysis in the NOWAC study. It is a prospective population-based cohort that tracks 34% (170.000) of all Norwegian women born between 1943 and 1957 [11]. We started the data collection in 1991 with surveys that cover topics including: the use of oral contraceptives and hormonal replacement therapy, reproductive history, smoking, physical activity, breast cancer, and breast cancer in the family. We also periodically update the study with data from The Norwegian Cancer Registry, and the Cause of Death Registry. In addition to the questionnaire data, we collected blood samples from 50.000 women, as well as more than 300 biopsies. From the biological samples we generated the first microarray based gene expression dataset in 2009, and later miRNA, DNA methylation, metabolomics, and RNA-seq datasets.

The data in the NOWAC cohort allows for a number of different study designs. While it is a prospective cohort study, we can also draw a case-control study from the cohort, or a cross-sectional study. We have published papers analyzing the questionnaire data (e.g. [14], [15]), and many research papers that investigate the questionnaire data together with the gene expression datasets (e.g. [16], [17]). We have also used the gene expression datasets to explore gene expression signals in blood and interactions between the tumor and the blood systemic response of breast cancer patients [18], [19]. Some analyses have resulted in patents [20] and commercialization efforts. There are still, however, many unexplored areas in the NOWAC datasets.

In the NOWAC study we are a group of researchers, PhD students, post docs, technical staff, and administrative staff. The researchers have backgrounds from statistics, medicine, epidemiology, and computer science. The administrative and the technical staff are responsible for managing the data, both data collection and data delivery to researchers. The interdisciplinary work and the complexity of the studies makes data management and analysis especially challenging.

### Data management and analysis

Surveys are the traditional data collection method in epidemiology. But today, questionnaire responses are increasingly integrated with molecular data. However, surveys are still important for designing a study that can answer particular research questions. In this section we describe how such omics data analysis was done in NOWAC before we developed our approach. We believe many studies have been, or are still, analyzing epidemiological data using a similar practice, and that our approach and lessons learned presented here will be useful for these.

In the NOWAC study we have stored the raw survey and registry data in an in-house database. Researchers apply to get questionnaire data variables exported from the database by scientific staff. This was typically done through SAS scripts that did some preprocessing, e.g. selecting applicable variables or samples, before the data was sent to researchers as SAS data files. The downstream analysis was typically done in SAS. Researchers used e-mail to communicate and send data analysis scripts, so there was no central hub with all the scripts and data.

In addition to the questionnaire data, the NOWAC study also integrates with registries (cancer and death) that are updated regularly. The datasets received from the different registries are typically delivered as comma-separated values (CSV) files to our scientific staff, which are then processed into a standardized format. Since the NOWAC study is a prospective cohort, some women are expected to get a cancer and move from the list of controls into the list of cases. This also requires updating their status in the analyses using gene expression data, and it makes it necessary to keep track of the case-control changes.

In the NOWAC study, we have analyzed our biological samples in labs outside our research institution. The received raw instrument datasets are then stored on a local server and made available to researchers on demand. Because of the complexity of the biological datasets, many of these require extensive preprocessing before they are ready for analysis.

### Issues in previous practice

Through nearly a decade of experiences from transcriptomics data analysis, we identified a set of issues with our previous practice that prevented us from fully ensuring reproducible data analysis:

1. It was difficult to keep track of the available datasets, how they were combined, and to determine how these had been processed. We had no standard data storage platform or structure, and there were limited reports for exported datasets used in different research projects.
2. There was no standard approach to preprocess and initiate data analysis. This was because the different datasets were analyzed by different researchers at different points in time, and there was little practice for sharing reusable code between projects.
3. It became difficult to reproduce the results reported in our published research manuscripts. This was because the lack of standardized preprocessing, sharing of analysis tools in their various versions, and full documentation of the analysis process.

## Enabling reproducible data analyses

To solve the above issues and enable easily reproducible research in the NOWAC study, we developed a system for managing and documenting the available datasets, a standardized data preprocessing and preparation system, and a set of best practices for data analysis and management. We first identified a set of requirements for a system to manage and document the different datasets:

1. It should provide users with a single interface to access the datasets, their respective documentation, and utility functions to access the raw and preprocessed data.
2. It should be capable of handling datasets in the order of a few GB and simultaneously retain interactive computation time for the analyses.
3. It should provide version history for the data and analysis code and tools.
4. The system should provide reproducible data analysis reports for modified datasets.
5. It should be portable and reusable by other systems or applications.
6. The system should be able to handle access management, data protection, and privacy concerns, such as anonymization.

To satisfy the above requirements we developed the *NOWAC R package*, a software package in the R programming language to provide access to all data, documentation, and utility functions.

We identified a set of requirements for this data preprocessing and preparation system as well:

1. The data preprocessing and preparation system should provide users with an interactive point-and-click interface to generate analysis-ready datasets from the NOWAC study.
2. It should use the *NOWAC R package* to retrieve datasets.
3. It should provide users with a list of possible options for filtering, normalization, and other options required to preprocess a microarray dataset.
4. It should generate a reproducible report along with any exported dataset.
5. It should export the data in a format that encourages following best practices for reproducible research in further downstream analyses.

Finally, we developed a set of best practices for data analysis in our study. In the remainder of the section we detail how we built the *NOWAC package*, the *Pippeline*, and discuss best practices for data analysis.

### The NOWAC R package: data management

The *NOWAC R package* is our solution for storing, documenting, and providing utility functions to parse and process the raw omics data in the NOWAC study (Figure 1). We use *git* to version control both the analysis code and datasets, and store the repository on a self-hosted git server. We bundle together all datasets in the *NOWAC package*. This includes both questionnaire, registry, and gene expression datasets. Because these are small by modern standards (currently all dataset are less than 10 GB) we are able to distribute them with our R package. Some datasets require preprocessing and quality control steps such as the removal of observations marred by technical artefacts (we sometimes refer to this as outlier removal) before the analysts explores the datasets. For this, we store the raw datasets, and the results of quality assessment. We store links to the raw datasets in their original file format, and as R data files to simplify importing in R. In addition, we store the R code we used to generate the R objects. For clarity, we decorate the scripts with specially formatted comments that can be used with *knitr* [21] to automatically generate data analysis reports. The reports highlight the transformation of the data from raw to processed and detail all information necessary to reproduce the entire processing, such as the specification of removed samples.

**Figure 1:**
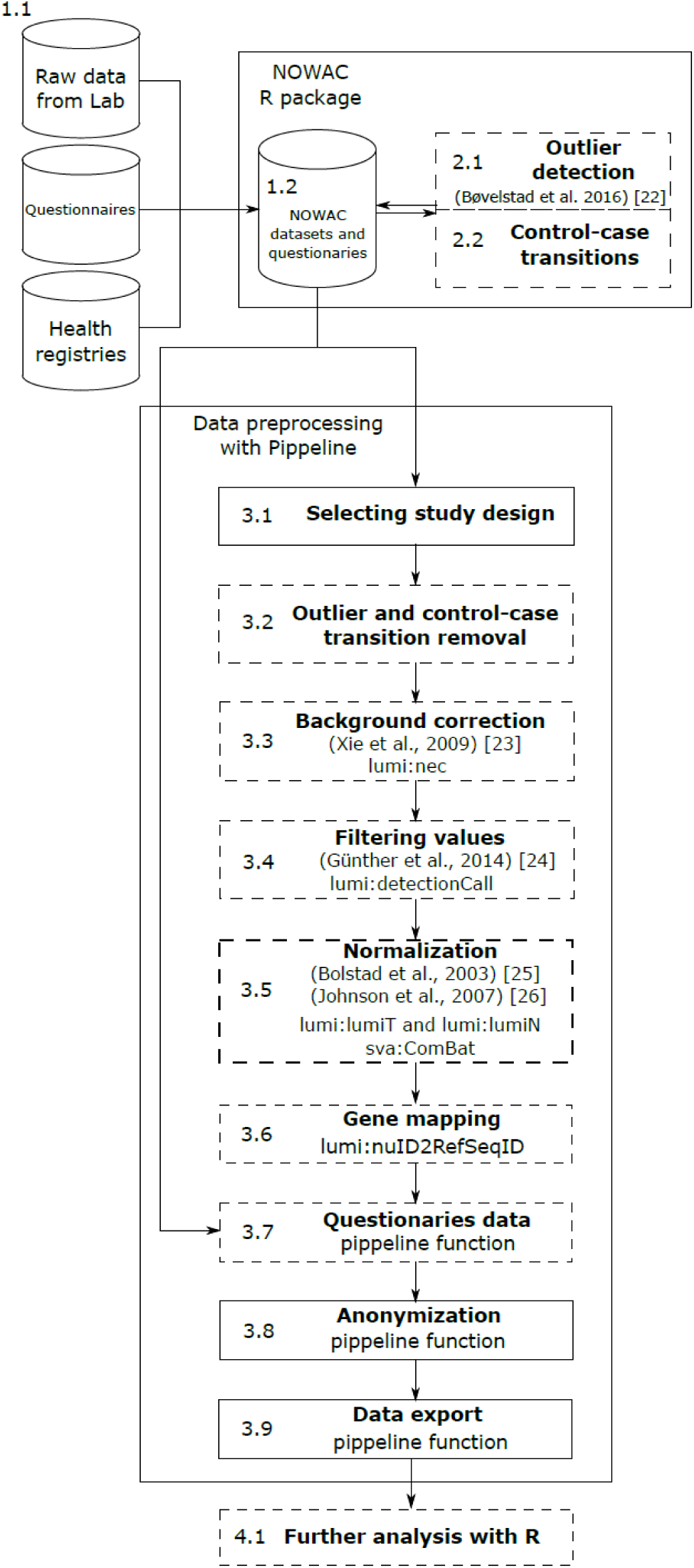
The standardized data processing pipeline for gene expression data preprocessing in the NOWAC study. Steps with a dashed border are optional, while steps with a solid border are mandatory. More details are in [22]–[26]

We have documented every raw dataset in the *NOWAC R package*. The documentation includes information such as data collection date, instrument types, the persons involved with data collection and analysis, and pre-processing methods. When users install the *NOWAC R package* the documentation is used to generate interactive help pages which they can browse in R, either through the command line or through an integrated development environment (IDE), such as RStudio. We can also export this documentation to a range of different formats, and researchers can also view them in the RStudio interface. Figure 2 shows the user interface of RStudio where the user has opened the documentation page for one of the gene expression datasets.

**Figure 2:**
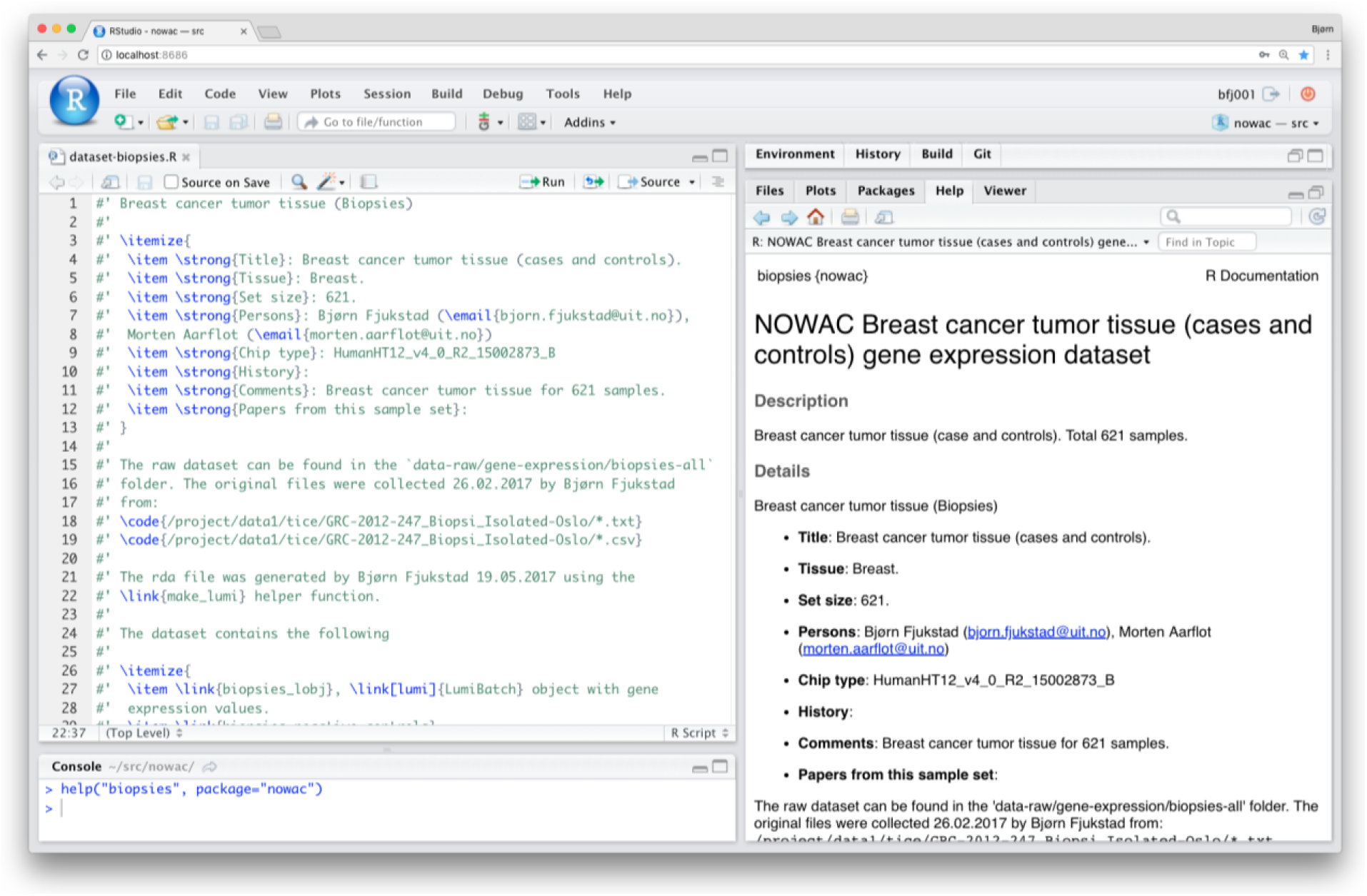
A screenshot of the user interface in R Studio, viewing the documentation help page for the “Biopsies” dataset in the NOWAC study. The right-hand panel shows the documentation generated by the code in the top left panel. The bottom left panel shows the R command that brought up the help page.

We use a single repository for the R package and put each dataset into a git submodule (Figure 3). This allows us to separate access to the datasets from the documentation and analysis code for data security and privacy reasons. Everyone with access to the repository can view the documentation and analysis code, but only a few have access to the data. Submodules allow us to keep the main repository size small, while still versioning the data. The *NOWAC R package* also provides various utility functions to process the raw datasets, and helper functions to retrieve questionnaire data.

**Figure 3:**
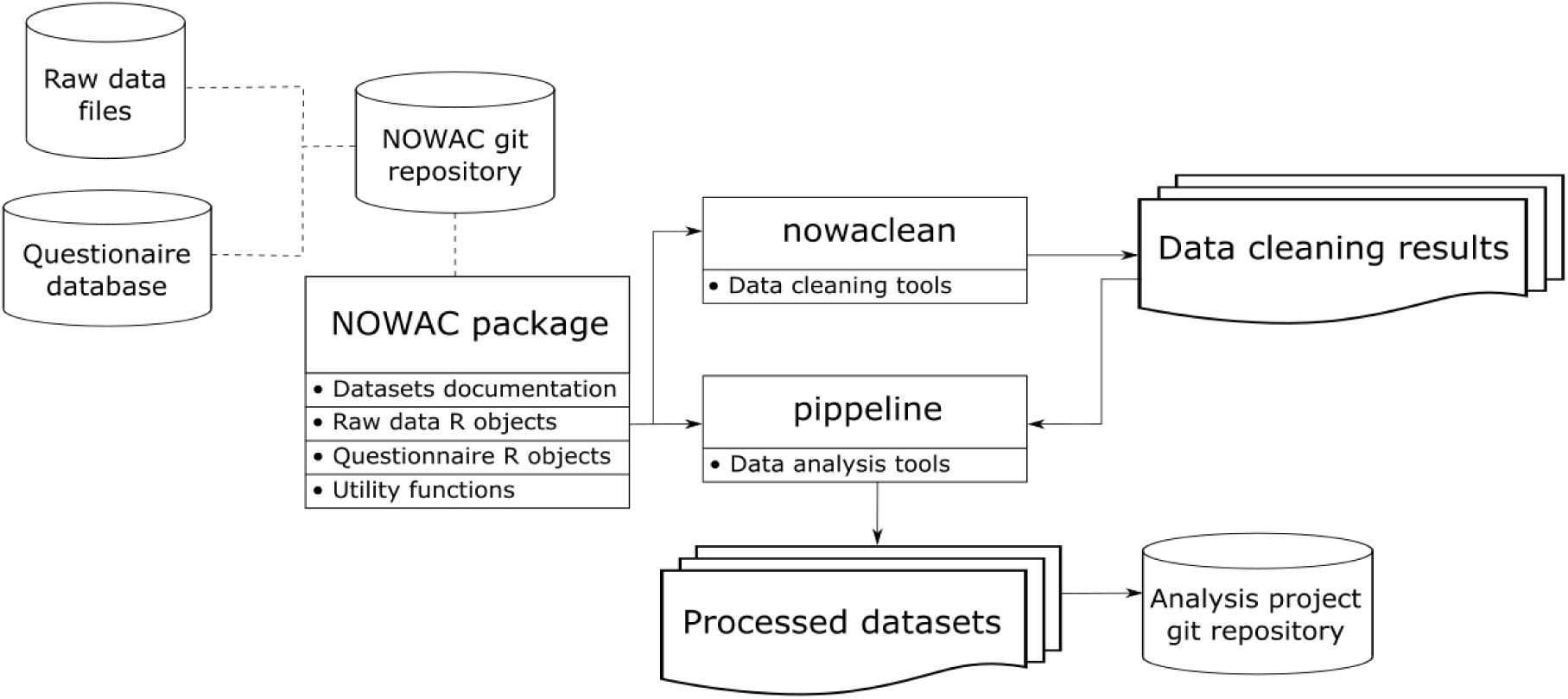
NOWAC R package and Pippeline deployment.

### Pippeline: Interactive Preprocessing Web Application

The use of the biological data in the NOWAC study in a research project comprise four steps (Figure 1). First, as explained above, the raw datasets are added to the *NOWAC R package* and documented thoroughly by a data manager. Second, we do manual quality assessment of the biological datasets. We add information about technical outliers to the *NOWAC R package* along with reports that describe why an observation is marked as an outlier. Third, the data manager generates an analysis-ready gene expression dataset for subsequent analysis using the interactive *Pippeline* tool as described below. Fourth, researchers further analyse the exported dataset with their tools of choice, following best practices for reproducible data analysis.

We have developed our preprocessing pipeline for gene expression data as a point-and-click web application called *Pippeline. Pippeline* generates an analysis-ready dataset by integrating biological datasets with questionnaire and registry data, all found in our *NOWAC package*. It allows selecting study design, removing already-discovered technical outliers, data normalization methods, filter values, and questionnaire fields. It presents the user with a list of possible processing options. We provide summary statistics for samples and probes about the changes made on each processing step in real time, so *Pippeline* users can see how each preprocessing step changes the number of samples and probes in the dataset (Figure 4). *Pippeline* exports the analysis-ready R data files, an R script that has all the choices and selections made during the preprocessing, and an R markdown which contains a human-readable report that can be used in the Methods section of a paper. The R script enables reproducing the output and intermediate data if needed.

**Figure 4:**
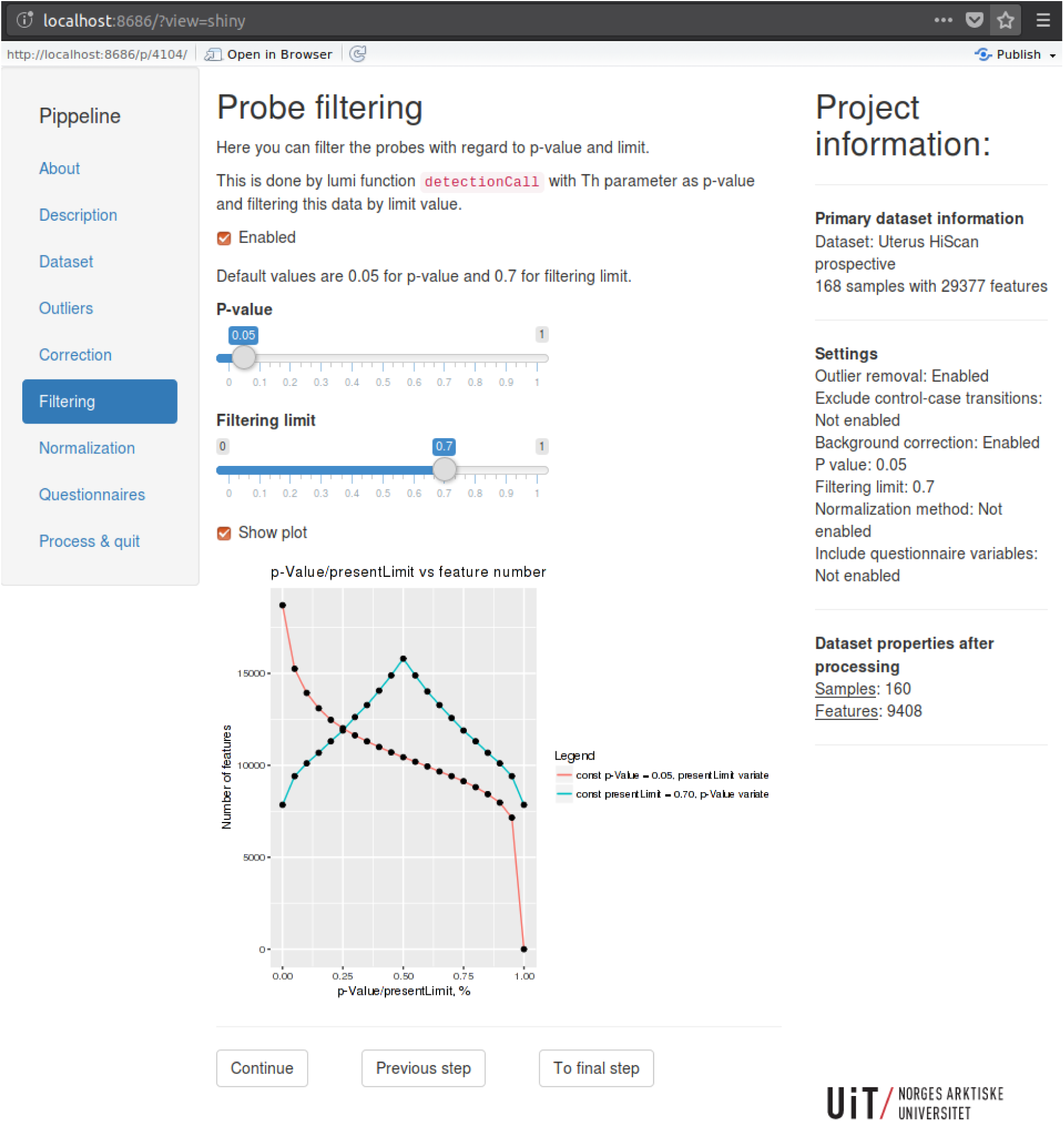
*A screenshot of the web-interface of Pippeline*. In the filtering step users specify the p-value and filtering limit for excluding gene expression probes in the dataset.

*Pippeline* is implemented in R as a web application using the Shiny framework [27]. It uses the *NOWAC R package* to retrieve all datasets. Intermediate data processing is implemented by creating temporary R files with the necessary functions for steps that are executed when the user interacts with the web application. Since our microarray datasets are small, the processing is very fast. Usually the *Pippeline* processing takes about 40 seconds. In the final step of the *Pippeline* we create a git repository with the output files, clone the repository to a folder on the user’s home directory, and create an RStudio project.

### Study-specific data analysis and result interpretation using R

Study-specific analyses are done by researchers using their methods and tools of choice, for example in RStudio. To encourage best practices for reproducible research we provide the following measures:

First, *Pippeline* exports the data as an RStudio project file with the data stored in a git repository. RStudio provides a graphical user interface for using git to version the code and data. This makes it easy to start using version control for any researcher.

Second, the *NOWAC R package* provides documentation about datasets. Missing information and corrections can be added either as suggested changes to the package or as issues to the package repository. This makes it easy to keep the documentation up to date.

Third, we provide a server with the necessary computational resources and software for the analysis, and we do not allow the data to be copied to another system. This makes it easy to keep all the data and code in one system, and to employ proper access management.

Finally, we encourage using best-practice regarding software and data management in our research group, and we give tutorials and workshops to teach these practices.

### Deployment of the NOWAC R package and Pippeline

We have deployed the *NOWAC R package* and *Pippeline* on two machines, and in addition we use our university’s storage system for raw data storage, and a database server for the questionnaire data. The storage machine runs the git and git-lfs servers used by the *NOWAC package*, and by the individual research projects. Only a few selected users have access to this machine. Another computer is used by the NOWAC researchers for their study-specific analyses to run the *Pippeline*. This machine has an RStudio server that the user can access through the browser. The machine also has home directories for the research projects. Finally, the researchers have their own laptops and workstations, used solely to establish a connection to the servers. No data should be copied out of the servers.

### Datasets stored in NOWAC R package and processed by Pippeline

Currently, we have used the *NOWAC package* and *Pippeline* for our 11 microarray datasets, but we are in the process of adding other data types also including microRNA, targeted RNA-seq, and methylation. The storage usage for the NOWAC package is 1.6 Gbytes including all R data objects. The total *Pippeline* output is 917 MBytes. The raw microarray (text) files are 8.7 GBytes in size, but the corresponding R objects are more efficiently stored.

### Best practices for reproducible epidemiological data analysis

From our experiences we have developed a set of best practices for data analysis. These apply both to researchers, developers, and the technical staff managing the data in a research study:

#### Document every step in the analysis

Analysis of modern datasets is a complex exercise with the possibility of introducing errors in every step. Analysts often use different tools and systems that require a particular set of input parameters to produce results. Thoroughly document every step from raw data to the final tables that go into a manuscript.

In the NOWAC study, we write help pages and reports for all datasets, and the optional pre-processing steps.

#### Generate reports and papers using code

With tools such as R Markdown [28] and *knitr* there are few reasons for decoupling analysis code with the presentation of the results through reports or scientific papers. Doing so ensures the correctness of reported results from the analyses, and greatly simplifies reproducing the results in a scientific paper.

In the NOWAC study we produce reports from R code. These include pre-processing and data delivery of datasets to researchers.

#### Version control everything

Both code and data changes over the course of a research project. Version control everything to make it possible to retrace changes and the person responsible for them. It is often necessary to roll back to previous versions of a dataset or analysis code, or to identify the researches that worked on specific analyses.

In the NOWAC study we encourage the use of git to version control both source code and data.

#### Collaborate and share code through source code management (SCM) systems

Traditional communication through e-mail makes it difficult to keep track of existing analyses and their design choices both for existing project members and new researchers. With SCM hosting systems such as Github, developing analysis code becomes more transparent to other collaborators, and encourages collaboration. It also simplifies the process of archiving development decisions such as choosing a normalization method.

In the NOWAC study we collaborate on data analysis through a self-hosted Gitlab [29] installation. We believe the ready-made git repository output from *Pippeline* encourages good software development practices and provides a good foundation for effective collaborative work.

### Limitations

A potential drawback of using an R package that is version controlled in git to manage, document, and analyze research datasets for researchers is the prerequisite programming skills. While the topic of software engineering best practices may be absent in current research training of many researchers, we believe such skills will become increasingly common in the scientific community.

One possible limitation of our *NOWAC R package* is its size. Microarray datasets are relatively small compared to sequencing data, so new datasets may require using extensions to git, such as git-lfs as we used in walrus [9]. This may become necessary, since we are currently expanding the *NOWAC package* and creating interactive pipelines similar to the *Pippeline* workflow for RNA-seq, Methylation, and microRNA datasets.

## Conclusions

We have proposed an approach to enable reproducible analyses for epidemiological omics data analyses. Our solution consists of several software tools embedded in the proper methodology, as well as best practices, and it solves a number of challenges previously encountered in omics studies. Among the advantages of our approach are the proper separation of datasets and tools, access management, anonymization, tracking of software version and dataset changes, documentation of processing steps and corresponding parameters, as well as cross-platform support, an easy-to-use graphical interface, and low latency. While we have applied our approach to a specific epidemiological research study for successful verification, we believe that it is generalizable to other biomedical analyses and even other scientific disciplines.

The *NOWAC R package*, without our data and data documentation is available at: https://github.com/uit-bdps/nowaclite

*Pippeline* and a description of our microarray preprocessing pipeline are available at: https://github.com/uit-bdps/pippeline

A demo dataset from [22] is available at: https://doi.org/10.18710/FGVLKS

## Acknowledgements

The NOWAC study was supported by a grant from the European Research Council (ERC-AdG 232997 TICE). The funders had no role in the design of the study; in the collection, analyses and interpretation of the data; in the writing of the manuscript; or in the decision to submit for publication.

Some of the data in this article are from the Cancer Registry of Norway. The Cancer Registry of Norway is not responsible for the analysis or interpretation of the data presented.

Microarray service was provided by the Genomics Core Facility, Norwegian University of Science and technology, and NMC – a national technology platform supported by the functional genomics program (FUGE) of the Research Council of Norway.

## References

[1] “Reality check on reproducibility,” Nat. News, vol. 533, no. 7604, p. 437, May 2016.

[2] E. Lund and V. Dumeaux, “Systems epidemiology in cancer,” Cancer Epidemiol. Biomark. Prev. Publ. Am. Assoc. Cancer Res. Cosponsored Am. Soc. Prev. Oncol., vol. 17, no. 11, pp. 2954–2957, Nov. 2008.

[3] V. Gallo et al., “STrengthening the reporting of OBservational studies in Epidemiology-Molecular Epidemiology (STROBE-ME): an extension of the STROBE statement,” Eur. J. Epidemiol., vol. 26, no. 10, pp. 797–810, Oct. 2011.

[4] P. Ivie and D. Thain, “Reproducibility in Scientific Computing,” ACM Comput Surv, vol. 51, no. 3, pp. 63:1–63:36, Jul. 2018.

[5] P. Amstutz et al., “Common Workflow Language, v1.0.” 08-Jul-2016.

[6] E. Afgan et al., “The Galaxy platform for accessible, reproducible and collaborative biomedical analyses: 2016 update,” Nucleic Acids Res., vol. 44, no. W1, pp. W3–W10, Jul. 2016.

[7] J. Köster and S. Rahmann, “Snakemake—a scalable bioinformatics workflow engine,” Bioinformatics, vol. 28, no. 19, pp. 2520–2522, Oct. 2012.

[8] M. Zaharia et al., “Apache Spark: a unified engine for big data processing,” Commun. ACM, vol. 59, no. 11, pp. 56–65, Oct. 2016.

[9] B. Fjukstad, V. Dumeaux, M. Hallett, and L. Bongo, “Reproducible Data Analysis Pipelines for Precision Medicine,” bioRxiv, p. 354811, Jun. 2018.

[10] “Pachyderm - Scalable, Reproducible Data Science.” [Online]. Available: https://www.pachyderm.io/. [Accessed: 13-May-2019].

[11] E. Lund et al., “Cohort Profile: The Norwegian Women and Cancer Study—NOWAC—Kvinner og kreft,” Int. J. Epidemiol., vol. 37, no. 1, pp. 36–41, Feb. 2008.

[12] R. C. Gentleman et al., “Bioconductor: open software development for computational biology and bioinformatics,” Genome Biol., vol. 5, no. 10, p. R80, 2004.

[13] “The Comprehensive R Archive Network.” [Online]. Available: https://cran.r-project.org/. [Accessed: 13-May-2019].

[14] M. Busund, N. S. Bugge, T. Braaten, M. Waaseth, C. Rylander, and E. Lund, “Progestin-only and combined oral contraceptives and receptor-defined premenopausal breast cancer risk: The Norwegian Women and Cancer Study,” Int. J. Cancer, vol. 142, no. 11, pp. 2293–2302, 2018.

[15] I. T. Gram, M. A. Little, E. Lund, and T. Braaten, “The fraction of breast cancer attributable to smoking: The Norwegian women and cancer study 1991–2012,” Br. J. Cancer, vol. 115, no. 5, pp. 616–623, Aug. 2016.

[16] K. S. Olsen, C. Fenton, L. Frøyland, M. Waaseth, R. H. Paulssen, and E. Lund, “Plasma fatty acid ratios affect blood gene expression profiles--a cross-sectional study of the Norwegian Women and Cancer Post-Genome Cohort,” PloS One, vol. 8, no. 6, p. e67270, 2013.

[17] V. Dumeaux, K. S. Olsen, G. Nuel, R. H. Paulssen, A.-L. Børresen-Dale, and E. Lund, “Deciphering normal blood gene expression variation--The NOWAC postgenome study,” PLoS Genet., vol. 6, no. 3, p. e1000873, Mar. 2010.

[18] M. Holden, L. Holden, K. S. Olsen, and E. Lund, “Local in Time Statistics for detecting weak gene expression signals in blood – illustrated for prediction of metastases in breast cancer in the NOWAC Post-genome Cohort,” Advances in Genomics and Genetics, 10-Jul-2017. [Online]. Available: https://www.dovepress.com/local-in-time-statistics-for-detecting-weak-gene-expression-signals-in-peer-reviewed-article-AGG. [Accessed: 27-Feb-2019].

[19] Vanessa Dumeaux et al., “Interactions between the tumor and the blood systemic response of breast cancer patients,” PLOS Comput. Biol., vol. to appear.

[20] V. Dumeaux and E. Lund, “Gene expression profile in diagnostics,” WO2014081313 A1, 30-May-2014.

[21] “Dynamic Documents with R and knitr,” CRC Press. [Online]. Available: https://www.crcpress.com/Dynamic-Documents-with-R-and-knitr/Xie/p/book/9781498716963. [Accessed: 27-Feb-2019].

[22] H. M. Bøvelstad, E. Holsbø, L. A. Bongo, and E. Lund, “A Standard Operating Procedure For Outlier Removal In Large-Sample Epidemiological Transcriptomics Datasets,” bioRxiv, p. 144519, May 2017.

[23] Y. Xie, X. Wang, and M. Story, “Statistical methods of background correction for Illumina BeadArray data,” Bioinformatics, vol. 25, no. 6, pp. 751–757, Mar. 2009.

[24] C.-C. Günther, M. Holden, and L. Holden, “Preprocessing of gene-expression data related to breast cancer diagnosis.” Norsk Regnesentral, 2014.

[25] B. M. Bolstad, R. A. Irizarry, M. Åstrand, and T. P. Speed, “A comparison of normalization methods for high density oligonucleotide array data based on variance and bias,” Bioinformatics, vol. 19, no. 19, pp. 185–193, Jan. 2003.

[26] W. E. Johnson, C. Li, and A. Rabinovic, “Adjusting batch effects in microarray expression data using empirical Bayes methods,” Biostatistics, vol. 8, no. 8, pp. 118–127, Jan. 2007.

[27] “Shiny.” [Online]. Available: https://shiny.rstudio.com/. [Accessed: 13-May-2019].

[28] “R Markdown.” [Online]. Available: https://rmarkdown.rstudio.com/. [Accessed: 13-May-2019].

[29] “The first single application for the entire DevOps lifecycle,” GitLab. [Online]. Available: https://about.gitlab.com/. [Accessed: 13-May-2019].

